# Optimal protein production by a synthetic microbial consortium: Coexistence, distribution of labor, and syntrophy

**DOI:** 10.1101/2022.01.12.476046

**Authors:** Carlos Martínez, Eugenio Cinquemani, Hidde de Jong, Jean-Luc Gouzé

## Abstract

The bacterium *E. coli* is widely used to produce recombinant proteins such as growth hormone and insulin. One inconvenience with *E. coli* cultures is the secretion of acetate through overflow metabolism. Acetate inhibits cell growth and represents a carbon diversion, which results in several negative effects on protein production. One way to overcome this problem is the use of a synthetic consortium of two different *E. coli* strains, one producing recombinant proteins and one reducing the acetate concentration. In this paper, we study a chemostat model of such a synthetic community where both strains are allowed to produce recombinant proteins. We give necessary and sufficient conditions for the existence of a coexistence equilibrium and show that it is unique. Based on this equilibrium, we define a multi-objective optimization problem for the maximization of two important bioprocess performance metrics, process yield and productivity. Solving numerically this problem, we find the best available trade-offs between the metrics. Under optimal operation of the mixed community, both strains must produce the protein of interest, and not only one (distribution instead of division of labor). Moreover, in this regime acetate secretion by one strain is necessary for the survival of the other (syntrophy). The results thus illustrate how complex multi-level dynamics shape the optimal production of recombinant proteins by synthetic microbial consortia.

## 1 Introduction

*Escherichia coli* (*E. coli*) is one of the most widely used bacteria for large-scale production of recombinant proteins such as insulin or the human growth hormone (Baeshen et al., 2015; Huang et al., 2012). The preferred carbon source for *E. coli*, as for many other bacteria, is glucose, supporting a faster growth rate compared to other sugars (Görke and Stülke, 2008; Wolfe, 2005). One problem with *E. coli* cultures grown on glucose is that fast growth leads to the secretion of acetate, a phenomenon known as overflow metabolism (Basan et al., 2015; Enjalbert et al., 2017). The resulting accumulation of acetate in the medium inhibits growth and represents a diversion of carbon, thus resulting in several negative effects on protein production (Eiteman and Altman, 2006; Luli and Strohl, 1990).

Different strategies have been proposed to overcome acetate formation. For example, some strategies prevent acetate overflow by deleting genes in acetate metabolism or by forcing cells to take up glucose at a rate below the overflow threshold (De Mey et al., 2007; Eiteman and Altman, 2006). These strategies have several inconveniencies, including suboptimal growth or secretion of other fermentation products. Other strategies aim to remove acetate from the culture medium. Acetate removal can be done, for example, with a dialysis reactor (Fuchs et al., 2002) or with macroporous ion-exchange resins (Huang et al., 2012). A promising alternative, inspired by naturally occurring syntrophic microbial consortia (Rosenzweig et al., 1994), consists in introducing an additional *E. coli* strain which has been metabolically engineered to consume acetate (Bernstein et al., 2012).

Along these lines, in recent theoretical work Mauri et al. (2020) proposed a coarse-grained mathematical model of a syntrophic consortium and investigated under which conditions it could improve the production of recombinant proteins as compared to mono-cultures. The model accounts for two different *E. coli* strains growing together: one producing the protein of interest (producers), and one reducing the presence of acetate (cleaners). As highlighted by the authors, optimizing the production of recombinant proteins in such a mixed culture may lead to gains in productivity (gram product per hour), but at the expense of diverting subtrate away from the producer to the cleaner, thus leading to a lower process yield (gram product per gram substrate). This trade-off between productivity and yield on the consortium level is a generalization of a well-known trade-off on the level of individual species, where high yield of protein production comes with an increased metabolic load that limits growth and therefore productivity (Kurland and Dong, 1996; Wu et al., 2016). Optimizing the performance of microbial consortia is a challenge for current industrial applications (Hays et al., 2015; Jagmann and Philipp, 2014; Pandhal and Noirel, 2014; Roell et al., 2019), because it requires the simultaneous optimization of individual microbial species and their interactions (Shong et al., 2012). Mathematical modeling is of great help in understanding and optimizing such complex multi-level dynamical systems.

In this work, we propose a theoretical study on the coexistence and optimization of a modified version of the above synthetic producer-cleaner consortium in which both *E. coli* strains produce recombinant proteins. We hypothesize that this may alleviate the productivity-yield trade-off in that some of the carbon diverted to the cleaners can be utilized for protein production. We first propose an extension of the mathematical model of Mauri et al. (2020) to account for the production of recombinant proteins by cleaners. Then, we study conditions for a coexistence equilibrium. In general, the long-term coexistence of two populations in a chemostat is not guaranteed. When two populations compete for a growth-limiting substrate, only one population survives (Hardin, 1960). Recent work on syntrophic relationships, in which one species produces an inhibitor (a metabolic by-product) that serves as a nutrient for the other species, have shown that coexistence is possible (Harvey et al., 2014; Heßeler et al., 2006; Sari et al., 2012; Stump and Klausmeier, 2016). The mixed culture studied in this work accounts for competition and syntrophy, so that establishing coexistence is not trivial. We determine necessary and sufficient mathematical conditions for a coexistence equilibrium.

Based on this equilibrium, we define a multi-objective optimization problem (MOP) for maximizing productivity and process yield. The solution of the MOP provides a graphical description, known as Pareto optimal front, of the combinations of productivity and process yield such that one metric cannot be improved without degrading the other (Miettinen, 2012). We show that, when cleaners are allowed to produce proteins as well, the Pareto optimal front is pushed towards higher levels of productivity and process yield. That is, the division of labor between a strain producing recombinant proteins and another strain cleaning up acetate is suboptimal as compared to the distribution of the production task over the two strains. Moreover, in order to attain the required growth rates along the Pareto optimal front, the cleaners need to take up not only glucose but also acetate secreted by the producers. In other words, the syntrophic relationship between the two strains is a necessary property for optimal production.

Our paper is organized as follows. In Section 2, we describe the chemostat model of the synthetic microbial community. In Section 3, we state Theorem 3.6 that gives necessary and sufficient conditions for the existence of a unique coexistence steady state. In Section 4, we define the process yield and productivity associated with each coexistence steady state. Then, we study the solutions of the multi-objective optimization problem (MOP) of simultaneously maximizing the process yield and productivity. The paper includes four Appendices with proofs and details of the algorithms to numerically determine the coexistence equilibrium and the solutions of the MOP.

## 2 Model description

The description of the model of the *E. coli* community follows that given by Mauri et al. (2020). Consider a chemostat (see Figure 1) in which two different strains of *E. coli* grow. We will refer to these strains as producers (with concentration *B*_*p*_) and cleaners (with concentration *B*_*c*_). Both *E. coli* strains can grow by taking up glucose (with concentration *G*). The glucose uptake rates are denoted 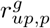 *and* 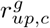 for producers and cleaners, respectively. The glucose uptake rate by producers is given by

**Figure 1:**
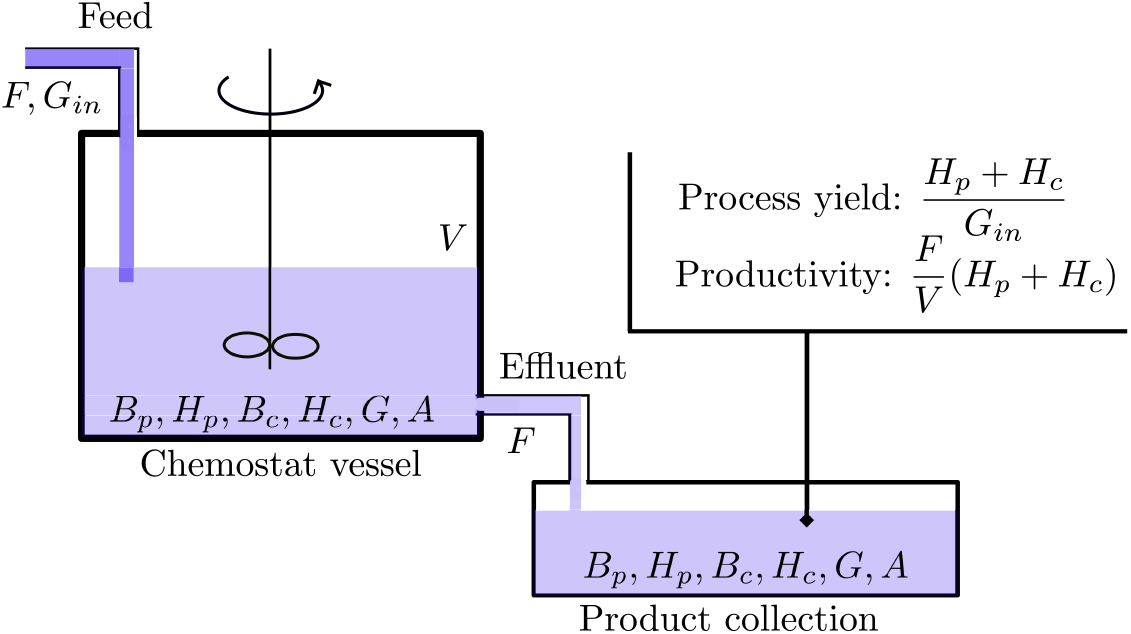
Schematic diagram of a chemostat used for the production of recombinant proteins (*H*_*p*_ + *H*_*c*_). The chemostat is fed at a rate *F* with a glucose concentration *G*_*in*_. The reactor is emptied at the rate *F* keeping a constant volume *V*. The concentrations of producers (*B*_*p*_) and cleaners (*B*_*c*_), glucose (*G*), and acetate (*A*) are homogeneous in the medium. The dilution rate *D* is defined as *F/V*. See Table 1 for the units of the variables.

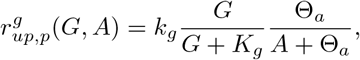

**Table 1:** Variables and rates in the model of producer-cleaner consortium given by the system (8)-(9).

| Variables | Unit | Description |
| --- | --- | --- |
| $B_p$ | gDW L<sup>-1</sup> | Biomass concentration of producers |
| $B_c$ | gDW L<sup>-1</sup> | Biomass concentration of cleaners |
| $G$ | g L<sup>-1</sup> | Concentration of glucose |
| $A$ | g L<sup>-1</sup> | Concentration of acetate |
| $H_p$ | gDW L<sup>-1</sup> | Concentration of recombinant proteins in producers |
| $H_c$ | gDW L<sup>-1</sup> | Concentration of recombinant proteins in cleaners |
| **Rates** |  |  |
| $r_{up,p}^g$ | h<sup>-1</sup> | Specific glucose uptake rate of producers |
| $r_{up,c}^g$ | h<sup>-1</sup> | Specific glucose uptake rate of cleaners |
| $r_{up,p}^a$ | h<sup>-1</sup> | Specific acetate uptake rate of producers |
| $r_{up,c}^a$ | h<sup>-1</sup> | Specific acetate uptake rate of cleaners |
| $r_{over,p}^a$ | h<sup>-1</sup> | Specific acetate secretion rate of producers |
| $r_{over,c}^a$ | h<sup>-1</sup> | Specific acetate secretion rate of cleaners |

where *k*_*g*_ is the maximal uptake rate of glucose, *K*_*g*_ is a half saturation constant, *A* is the acetate concentration, and Θ is a constant representing the inhibitory effect of acetate. In cleaners, the gene *ptsG* is deleted. This gene encodes a major subunit of the glucose uptake system (Deutscher et al., 2006), so that its deletion reduces the glucose uptake rate. 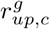 is then given by

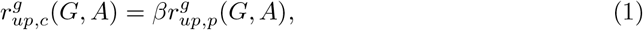

where *β* ∈ [0, 1] is a dimensionless constant, reflecting the deletion of the gene *ptsG*. If *β* = 1, the gene *ptsG* is present and both strains take up glucose at the same rate. The units of the variables and rates are summarized in Table 1.

When the glucose uptake rate of *E. coli* is above a threshold rate *l*, then cells secrete acetate through overflow metabolism (Basan et al., 2015). The overflow rates are denoted by 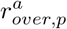 and 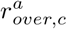 for producers and cleaners, respectively. They are defined as

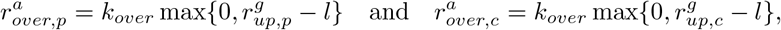

where *k*_*over*_ is a proportionality constant. Note that the maximal glucose uptake rate of cleaners is given by *βk*_*g*_. Thus, if

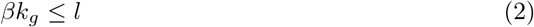

then cleaners cannot secrete acetate. This property is satisfied by the parameters in Table 2, in accordance with the fact that cleaners are primarily designed to remove acetate from the medium.

**Table 2:** Parameters and their values taken from the work of Mauri et al. (2020).

| Parameter | Value | Unit | Description |
| --- | --- | --- | --- |
| $k_g$ | 1.53 | $\text{h}^{-1}$ | Maximal glucose uptake rate |
| $K_g$ | 0.09 | $\text{g L}^{-1}$ | Glucose half-saturation constant |
| $\Theta_a$ | 0.52 | $\text{g/L}$ | Acetate inhibition constant |
| $k_a$ | 0.97 | $\text{h}^{-1}$ | Maximal acetate uptake rate |
| $K_a$ | 0.5 | $\text{g L}^{-1}$ | Acetate half-saturation constant |
| $\Theta_g$ | 0.25 | $\text{g L}^{-1}$ | Glucose uptake inhibition constant |
| $k_{over}$ | 0.17 | — | |
| $l$ | 0.7 | $\text{h}^{-1}$ | Overflow metabolism threshold rate |
| $Y_g$ | 0.44 | $\text{gDW g}^{-1}$ | Yield coefficient |
| $Y_a$ | 0.3 | $\text{gDW g}^{-1}$ | Yield coefficient |
| $\beta$ | 0.26 | — | |
| $k_{Acs}$ | 1.46 | $\text{h}^{-1}$ | |
| $K_{Acs}$ | 0.012 | $\text{g L}^{-1}$ | |
| $Y_{h/p}$ | 0 - 1 | | |
| $Y_{h/c}$ | 0 - 1 | | |
| $k_{deg}$ | 0.0044 | $\text{h}^{-1}$ | Degradation rate of producers |
| $D$ | - | $\text{h}^{-1}$ | Dilution rate |
| $G_{in}$ | - | $\text{g L}^{-1}$ | Input glucose concentration |

In the presence of glucose, *E. coli* cells cannot grow on acetate, a phenomenon known as Carbon Catabolite Repression (CCR) (Kremling et al., 2015; Wolfe, 2005). The uptake rates of acetate are denoted by 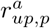 *and* 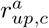 for producers and cleaners, respectively. The acetate uptake rate by producers is given by

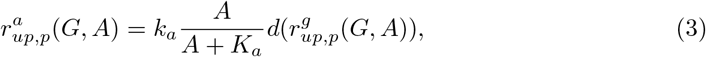

where *k*_*a*_ is the maximal acetate uptake rate, *K*_*a*_ is a half-saturation constant, and *d* is a down-regulation function accounting for CCR. Before describing the function *d*, we define 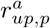. The assimilation of acetate by *E. coli* can be increased with a plasmid enabling the inducible overexpression of the native gene *acs*, coding for the enzyme acetyl-CoA synthetase (Lin et al., 2006). Hence, the uptake rate of acetate by cleaners is

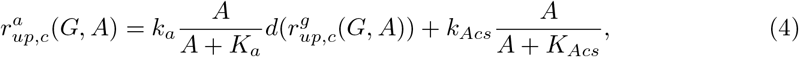

where *k*_*Acs*_ is the maximal acetate uptake rate due to overexpression of the gene *acs* and *K*_*Acs*_ is a half-saturation constant. Note that if cleaners are not genetically modified, then *β* = 1 (see (1)) and *k*_*Acs*_ = 0. In that case, both acetate and glucose uptake rates, are equal for both strains.

The down-regulation function *d* represents the inhibitory effect of glucose uptake and decreases as the glucose uptake rate increases. In the work of Mauri et al. (2020), the down-regulation function is defined as

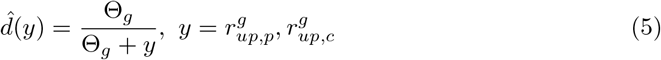

with Θ_*g*_ a constant. We modify the description of the down-regulation function (5) as follows:

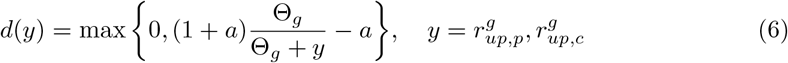

with *a* = Θ_*g*_*/l*. The main motivation to describe *d* as in (6) arises from the following property:

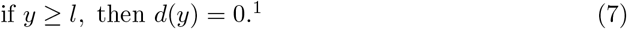

This property states that if overflow metabolism occurs (*i*.*e*., the glucose uptake rate is higher than *l*) then there is no acetate uptake, which is qualitatively consistent with CCR (Wolfe, 2005). An important advantage of (6) is that it facilitates the mathematical analysis of the model (see below). Figure 2 compares the two alternatives of *d*. In Appendix D, we show that our optimization results are insensitive to this model change.

**Figure 2:**
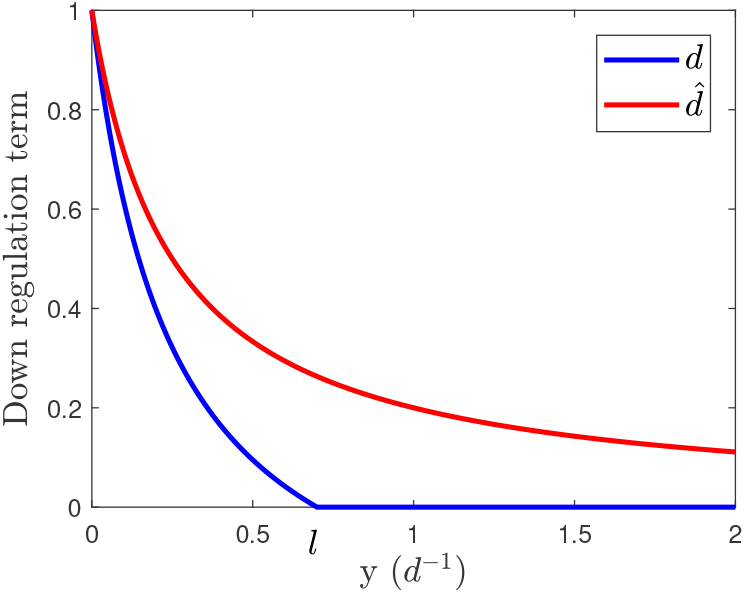
Down-regulation function *d* compared to the down-regulation function Θ_*g*_*/*(Θ_*g*_ + *y*). Parameters are taken from Table 2.

Both strains carry a plasmid for the expression of a recombinant protein. The concentration of proteins produced by producers and cleaners are denoted by *H*_*p*_ and *H*_*c*_, respectively. The fraction of biomass production assigned to the synthesis of recombinant protein is determined by the (dimensionless) product yield constants *Y*_*h/p*_ and *Y*_*h/c*_. In the model of Mauri et al. (2020), cleaners cannot produce heterologous proteins and they mainly serve to reduce the acetate concentration in the medium. In this new model, the carbon lost in the form of acetate can be recovered by cleaners and transformed into recombinant protein.

The specific rate of biomass production per unit of biomass, including *B*_*p*_ and *H*_*p*_, is given by

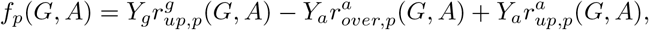

with *Y*_*a*_ and *Y*_*g*_ yield coefficients. Note that 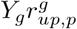 and 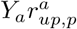 are gains of carbon due to glucose and acetate uptake, respectively, and 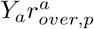 is a loss of carbon due to overflow metabolism. Similarly, the specific rate of cleaner biomass production per unit of biomass is given by

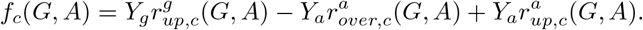

The evolution of *B*_*p*_, *B*_*c*_, *G*, and *A* is now defined by

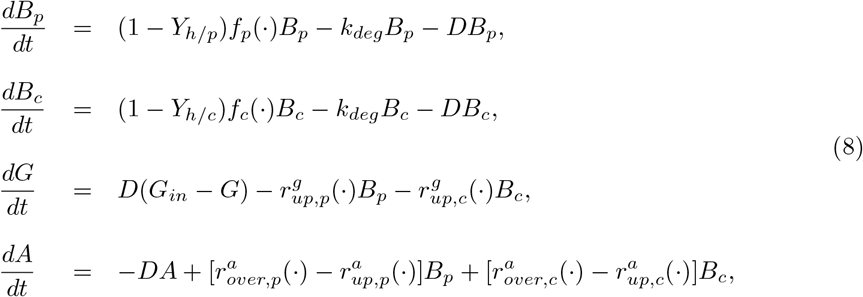

and the evolution of the recombinant protein concentrations is given by

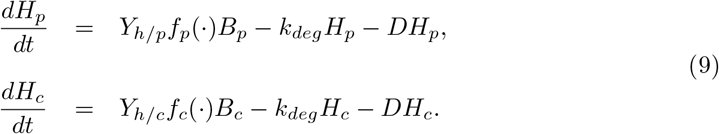

System (8)-(9) can be seen as an extension of the model of Mauri et al. (2020). Indeed, if *Y*_*h/c*_ = 0 and *d* is taken as (5), then (8)-(9) is equivalent to the model of Mauri et al. (2020). Note that the model has been intentionally split into two subsystems, (8) and (9), such that (8) is uncoupled from (9). To establish the existence of coexistence steady states, we only need to study system (8), but for optimizing the production of recombinant proteins, we must study both.

Figure 3 shows the steady-state solution of (8) obtained by simulation from three different kinds of initial conditions: the absence of cleaners, the absence of producers, the presence of both. These simulations suggest that (8) admits at most three non-trivial equilibria:

**Figure 3:**
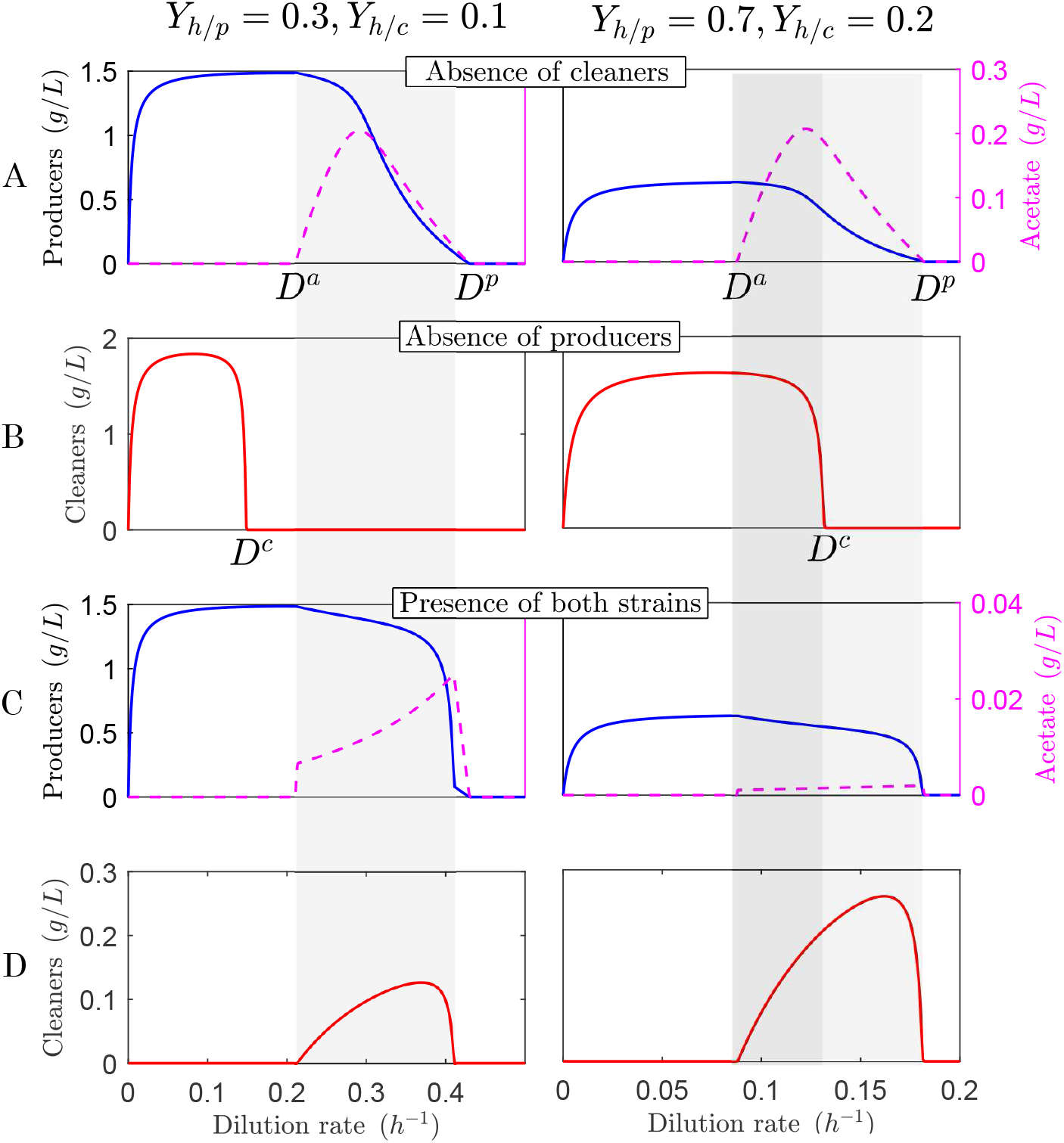
Steady-state solutions of (8) obtained by simulation for different values of the dilution rate and two different pairs of values for *Y*_*h/p*_ and *Y*_*h/c*_. *D*^*a*^ is the dilution rate above which acetate overflow occurs when producers grow individually. *D*^*p*^ and *D*^*c*^ are the dilution rates at which producers and cleaners go extinct when growing individually. The dilution rates *D*^*a*^, *D*^*p*^, and *D*^*c*^ are formally defined in Propositions 3.1 and 3.4. The shaded area represents the values of the dilution rate at which coexistence is possible. The darker shaded area corresponds to values of *D* lower than *D*^*c*^ and the lighter shaded area corresponds to values of *D* lower than *D*^*p*^ and higher than or equal to *D*^*c*^. **A**. *B*_*c*_(0) = 0, *B*_*p*_(0) *>* 0 (absence of cleaners). **B**. *B*_*c*_(0) *>* 0, *B*_*p*_(0) = 0 (absence of producers). **C and D**. *B*_*c*_(0), *B*_*p*_(0) *>* 0 (presence of producers and cleaners).

I. An equilibrium with the presence of producers but absence of cleaners (see Figure 3A). In this equilibrium the acetate concentration in the bioreactor may be zero or positive.
II. An equilibrium with the presence of cleaners but absence of producers. Since cleaners do not produce acetate (because (2) holds), the acetate concentration is zero.
III. A coexistence equilibrium (see Figure 3C and 3D).

Note that, at equilibrium, the growth rate of any species present in the bioreactor (that is, (1 − *Y*_*h/p*_)*f*_*p*_ − *k*_*deg*_ for the producer, and (1 − *Y*_*h/c*_)*f*_*c*_ − *k*_*deg*_ for the cleaner) must be equal to *D*. We are especially interested in coexistence equilibria. We observe the existence of an interval of values of *D* (shaded area), which depends on the values of *Y*_*h/p*_ and *Y*_*h/c*_, such that coexistence is possible. In the following section we determine necessary and sufficient conditions for the existence of a coexistence equilibrium.

## 3 Coexistence at steady states

In this section, we characterize the existence of coexistence equilibria of (8). We first present some results on the existence of equilibria with the presence of only one bacterial population. Then, based on these equilibria, we state the main result on coexistence. To state our results, some assumptions are necessary. The first assumption is

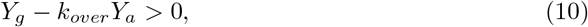

which is verified by the parameters in Table 2. The assumption implies that the function 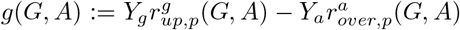 is strictly increasing with respect to *G* and strictly decreasing with respect to *A* (Martínez and Gouzé, 2021). This monotonicity of *g* is necessary to prove the uniqueness of the equilibrium with producers and no cleaners (Proposition 3.1). We also assume that

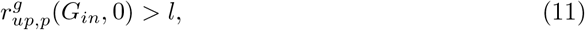

in agreement with the parameter values listed in Table 2. During long-term operation of the bioreactor in presence of bacteria, the glucose concentration in the medium cannot be higher than *G*_*in*_. Then, if (11) does not hold (*i*.*e*., 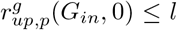), overflow metabolism is impossible in the long-term, and the dynamics of (8) is reduced to that of a classical chemostat model with two competitors for one substrate (glucose) (Smith and Waltman, 1995). Note that assumptions (10) and (11) have been used by Martínez and Gouzé (2021) to study a simplified version of (8) without cleaners.

The following proposition characterizes the existence of steady states without cleaners. The proof is adapted from Proposition 1 by Martínez and Gouzé (2021), where a similar result is presented, but in the absence of acetate consumption 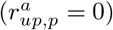.

### Proposition 3.1 (Producer steady state).

*Assume that (10) and (11) hold. Let us define D*^*p*^ := (1 − *Y*_*h/p*_)*f*_*p*_(*G*_*in*_, 0) − *k*_*deg*_. *We have:*

a. *If D*^*p*^ *> D, then (8) admits a unique equilibrium with the presence of producers and absence of cleaners, denoted by* 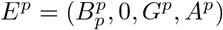. *Moreover, for D*^*a*^ = (1 − *Y*_*h/p*_)*Y*_*g*_*l* − *k*_*deg*_ *we have:*
  I. *If D*^*a*^ *> D, then A*^*p*^ = 0 *and* 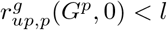.
  II. *If D*^*a*^ = *D, then A*^*p*^ = 0 *and* 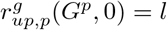.
  III. *If D*^*a*^ *< D, then A*^*p*^ *>* 0 *and* 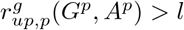.
b. *If D*^*p*^ ≤ *D, then (8) has no equilibrium with the presence of producers and absence of cleaners*.

*Proof*. See Appendix A.

### Remark 3.2 (Acetate secretion).

Proposition 3.1 shows the existence of a threshold dilution rate characterizing the presence of acetate in the medium, denoted by *D*^*a*^ (see Figure 3A). As we will see later, a coexistence equilibrium for (8) is only possible if the dilution rate is higher than *D*^*a*^. Proposition 3.1 also introduces *D*^*p*^, the threshold dilution rate above which producers grow extinct. No coexistence equilibrium is possible for dilution rates higher than *D*^*p*^.

### Remark 3.3 (Acetate consumption).

Note that there is no acetate consumption by the bacteria at equilibrium (the acetate secreted, if any, is diluted out at the same rate). Indeed, if *D* ≤ *D*^*a*^, there is no acetate, so there is trivially no acetate consumption. Conversely, if *D > D*^*a*^, there is overflow metabolism. Then, using (7), we have that there is no acetate consumption. The term 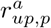 has therefore no influence on the steady state of producers. Thus, the non-trivial equilibrium determined by Martínez and Gouzé (2021) is exactly the same as the non-trivial equilibrium given by Proposition 3.1, even if the authors do not consider acetate consumption.

To study the existence of equilibria with cleaners and no producers, we assume that (2) holds. As discussed in Section 2, this implies that

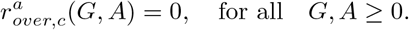

This means that cleaners cannot secrete acetate. This assumption is qualitatively consistent with the fact that cleaners are designed to mainly grow on acetate and is consistent with the parameters from Table 2.

### Proposition 3.4 (Cleaner steady state).

*Assume that (2) holds and let us define D*^*c*^ = (1 − *Y*_*h/c*_)*f*_*c*_(*G*_*in*_, 0) − *k*_*deg*_. *We have:*

a. *If D*^*c*^ *> D, then (8) admits a unique equilibrium with the presence of cleaners and absence of producers, denoted by* 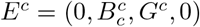.
b. *If D*^*c*^ ≤ *D, then (8) has no equilibrium with the presence of cleaners*.

*Proof*. See Appendix A.

### Remark 3.5 (Cleaner washout).

Proposition 3.4 introduces the threshold dilution rate *D*^*c*^ above which cleaners are washed out when grown without producers. As shown below, a coexistence equilibrium is possible at dilution rates higher than *D*^*c*^.

The main result of this section characterizes the existence of coexistence steady states for (8). This characterization is based on the positive steady states of the cleaners and producers growing separately.

### Theorem 3.6 (Coexistence steady states).

*Assume that (2), (10), and (11) hold. Let D*^*p*^ *and D*^*a*^ *be given by Proposition 3.1, and let D*^*c*^ *be given by Proposition 3.4*.

a. *If D* ≤ *D*^*a*^ *or D* ≥ *D*^*p*^, *then there is no coexistence equilibrium*.
b. *If D*^*a*^ *< D < D*^*p*^ *and D < D*^*c*^ *then a (unique) coexistence equilibrium is possible, if and only if*

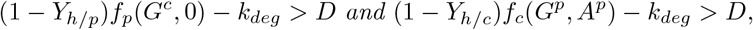

*with G*^*p*^ *and A*^*p*^ *defined in Proposition 3.1, and G*^*c*^ *defined in Proposition 3.4*.
c. *If D*^*a*^ *< D < D*^*p*^ *and D* ≥ *D*^*c*^ *then a (unique) coexistence equilibrium is possible, if and only if*

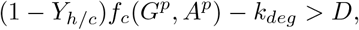

*with G*^*p*^ *and A*^*p*^ *defined in Proposition 3.1*.

*Proof*. See Appendix A.

### Remark 3.7 (Competition and syntrophy).

Note that the difference between cases (b) and (c) in Theorem 3.6 is the value of the dilution rate with respect to *D*^*c*^. If *D < D*^*c*^ (case (b)), according to Proposition 3.4, in the absence of producers, cleaners survive on glucose only and settle in an equilibrium. The presence of producers can be beneficial by providing acetate, which is known as protocooperation (Roell et al., 2019), or harmful through glucose competition. This can be observed in Figures 3B and 3D (right-hand side). When *D* is higher than but close to *D*^*a*^, cleaners reach a higher density when grown alone than when growing together with producers (competition). However, when *D* is lower than but close to *D*^*c*^, cleaners reach a lower density when grown alone than when growing together with producers (protocooperation). Now, if *D* ≥ *D*^*c*^ (case (c)), cleaners go extinct when growing alone. Therefore, if coexistence is possible, this is due to the syntrophic relationship in which the producers provide acetate to cleaners. As we will see later, synthrophy seems to play an important role in the performance of the system.

The following corollary gives a characterization of the existence of coexistence equilibria based on dynamical properties of the non-coexistence equilibria.

### Corollary 3.8.

*Assume that (10), (11), and (2) hold. Let E*^*p*^ *and E*^*c*^ *be the equilibria given by Propositions 3.1 and 3.4, respectively, whenever they exist. Let D*^*a*^ *be given by Proposition 3.1 and assume that D > D*^*a*^.

a. *If E*^*p*^ *does not exist, then there is no coexistence equilibrium*.
b. *If E*^*p*^ *exists and E*^*c*^ *does not exist, then there is a unique coexistence equilibrium if and only if E*^*p*^ *is hyperbolic and unstable*.
c. *If E*^*p*^ *and E*^*c*^ *exist, then there is a unique coexistence if and only if E*^*p*^ *and E*^*c*^ *are unstable and hyperbolic*.

*Proof*. The statement follows from applying Lemmas A.7 and A.8 in Appendix A.

### Remark 3.9 (Dynamics on the boundary).

According to Lemmas A.7 and A.8 in Appendix A, if *E*^*p*^ and *E*^*c*^ exist, the Jacobian matrices associated with them each have three negative eigenvalues. Therefore, the conditions in Corollary 3.8 state that the fourth eigenvalue must be negative. In such conditions, from a stable manifold argument, *E*^*p*^ and *E*^*c*^ can only be reached by solutions starting on 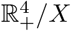, with *X* the set characterized by *B*_*c*_ *>* 0 and *B*_*p*_ *>* 0 (*i*.*e*., 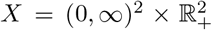). Thus, *E*^*p*^ and *E*^*c*^, whenever they exist, must repel solutions starting in *X* to ensure the existence of a coexistence equilibrium. This characterization can be compared with some classical results from the theory of persistence (Smith and Thieme, 2011), where the existence of a coexistence equilibrium can be determined from the dynamics on the boundary 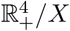 (*i*.*e*., the set where *B*_*c*_ = 0 or *B*_*p*_ = 0).

We end this section with a lemma stating how to determine the protein concentrations associated with the coexistence equilibrium given by Theorem 3.6.

### Lemma 3.10.

*Let* 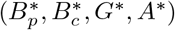 *be a coexistence equilibrium of (8). Then, the protein concentrations associated with this equilibrium are given by*

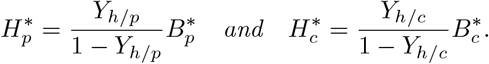

*Proof*. See Appendix A.

We conclude the section by observing that, for a fixed set of parameter values (*e*.*g*., those in Table 2), the stability of the above equilibria can be verified by the same numerical/algebraic methods utilized in Mauri et al. (2020).

## 4 Optimal performance at the coexistence equilibrium

The necessary and sufficient conditions given by Theorem 3.6 for the existence of a coexistence equilibrium allow for a proper study of the optimization of (8)-(9) at steady state. While the statement of Theorem 3.6 ensures the existence of a non-empty region where coexistence is possible, its proof indicates how to construct an efficient algorithm for determining numerically the coexistence equilibria (see Appendix B). In this section, we define a multiobjective optimization problem (MOP) aiming to maximize the process yield (g product per g substrate) and protein volumetric productivity g per L per h). The process yield is especially relevant in a biotechnological context when the selling price of the product is low, since the cost of glucose becomes a significant fraction of the value of the product. Volumetric productivity determines the rate at which the product can be formed, and thus dictates the overall volume needed for a given plant output (Van Dien, 2013).

The decision variables in the optimization problem are chosen to be the dilution rate (*D*) and the product yield of each strain (*Y*_*h/p*_ and *Y*_*h/c*_). The dilution rate is well-known to be an important operational variable that is optimal at intermediate values (Doran, 1995). Similarly, the choice of product yields is not trivial. Low values of *Y*_*h/p*_ naturally lead to low protein production, while high values lead to more recombinant proteins but poor cell growth resulting again in low productivity. Another operational parameter is the input glucose concentration *G*_*in*_. However, as we will show later, performance always improves with an increase of this parameter, making its choice trivial (see also the work of Mauri et al. (2020)).

In mathematical optimization, the feasible region corresponds to the set of all possible combinations of the decision variables. Since we are interested in coexistence equilibria, we consider the feasible region 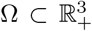, such that for any *v* := (*Y*_*h/p*_, *Y*_*h/c*_, *D*) ∈ Ω, (8) admits a coexistence equilibrium. The region Ω can be determined by the conditions given by Theorem 3.6 and it looks like a cone with a vertex close to (1, 1, 0) (see Figure 4). For any *v* ∈ Ω, we will denote by 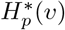 and 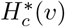 the protein concentrations associated with producers and cleaners, respectively, at the coexistence steady state (see Lemma 3.10). We define the objective function 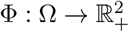 by

**Figure 4:**
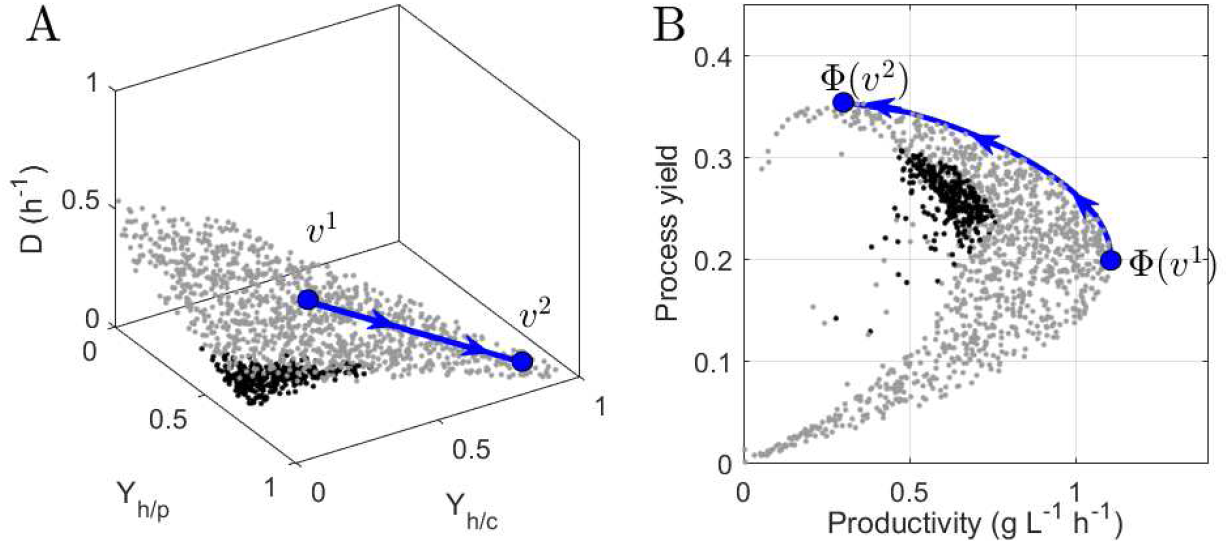
**A**. Region Ω represented by 3000 points. Gray points correspond to *D* ≥ *D*^*c*^ and black points correspond to *D < D*^*c*^ (see Remark 3.7). The point *v*^1^ = (0.51, 0.34, 0.28) maximizes productivity (Φ_1_) and *v*^2^ = (0.91, 0.88, 0.04) maximizes the process yield (Φ_2_). The blue curve is such that its image through Φ is the POF. **B**. Function Φ evaluated at the points represented in A. The blue curve is the POF.

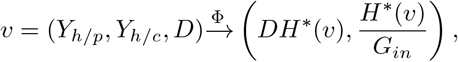

where 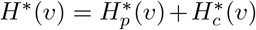. The quantities *DH*\*(*v*) and *H*\*(*v*)*/G*_*in*_ are the steady-state productivity and process yield, respectively. We want to solve the following MOP:

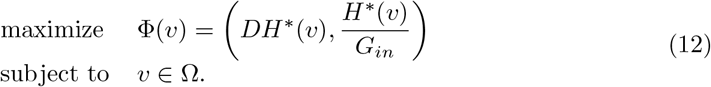

We look for Pareto optimal solutions, that is, solutions that cannot be improved in any of the objectives (process yield or productivity) without degrading the other objective. Generally, there is no single Pareto optimal solution optimizing both objectives, but a set of Pareto optimal solutions called the Pareto optimal front (POF). The structure of the sets Ω and Φ(Ω) plays an important role in the success of the employed numerical method to solve (12). Pareto curves cannot be computed efficiently in many cases, especially in the non-convex case where methods such as *ϵ*-constraint are necessary (Gunantara, 2018). Figure 4 reveals a favorable structure of Ω and Φ(Ω) (scatter plots) for applying the weighting method (Miettinen, 2012). Accordingly, the problem (12) is transformed into the following so-called weighting problem:

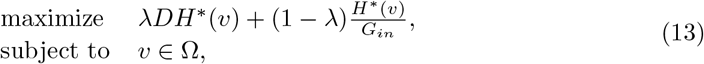

where *λ* ∈ [0, 1]. It is well known that if Ω and Φ(Ω) are convex, then *v*\* is a Pareto optimal solution if there is *λ* ∈ [0, 1] such that *v*\* is a solution to the weighting problem (13) (see Corollary 3.1.8 by Miettinen (2012)).

For any *λ* ∈ [0, 1], problem (13) is solved numerically with the interior point algorithm implemented in the toolbox *fmincon* of MATLAB (Byrd et al., 1999). Before using *fmincon*, some technical details must be addressed. For example, the feasible region must be a closed set and the objective function must be continuous on this set. For ease of reading, we discuss such technical details in Appendix C.

Figure 4 shows the POF obtained using the weighting method and the subset of Ω whose image is the POF. From this figure we observe that:

a. The weighting method accurately returns the POF.
b. The set of all points *v* ∈ Ω such that Φ(*v*) belongs to the POF can be approximated by the line joining the points *v*^1^ and *v*^2^, corresponding to maximum productivity and maximum yield, respectively.
c. Along the POF, the process yield decreases as the productivity increases.

Most importantly, regarding the syntrophy-competition relationship between both strains, the image Φ of all points (*Y*_*h/p*_, *Y*_*h/c*_, *D*) ∈ Ω such that *D < D*^*c*^ is a dominated solution (black points). In other words, if cleaners can survive when growing individually, then the system is operated under suboptimal conditions (see Remark 3.7).

### 4.1 Impact of *G*_*in*_

Figure 5 shows the effects of varying the input glucose concentration *G*_*in*_. According to Figure 5B, as *G*_*in*_ increases, the POF expands in such a way that the POF dominates the POFs associated with lower values of *G*_*in*_. This shows why *G*_*in*_ is not a relevant decision variable for the MOP. The input glucose concentration must always be chosen as high as possible. We note that the maximal process yield slightly increases (Φ_2_), while the maximal productivity increases almost linearly with *G*_*in*_ (Φ_1_). Figure 5A shows that the values of the decision variables associated with each POF are not much affected by the value of *G*_*in*_.

**Figure 5:**
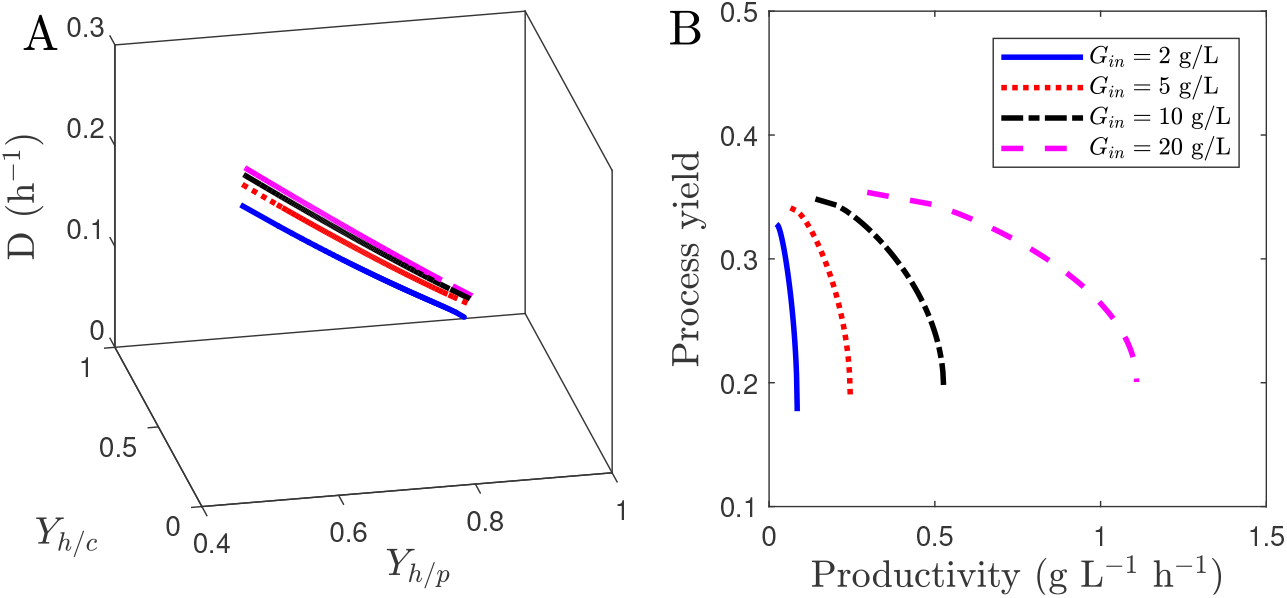
Influence of *G*_*in*_ on the POF. **A**. Decision variables which image through Φ is a POF. **B**. POF for different values of *G*_*in*_.

### 4.2 Comparison with the monoculture

The potential of a consortium over a monoculture, in terms of productivity, has been demonstrated experimentally by Bernstein et al. (2012) and theoretically by Mauri et al. (2020), but limited to the case that only producers synthesize the target protein. Figure 6A confirms those findings in the more general case of both species contributing to the synthesis process, showing that productivity can increase 63% in case of a consortium of producers and cleaners. However, it also shows that the monoculture can reach a higher process yield, although of course at the expense of low productivity.

**Figure 6:**
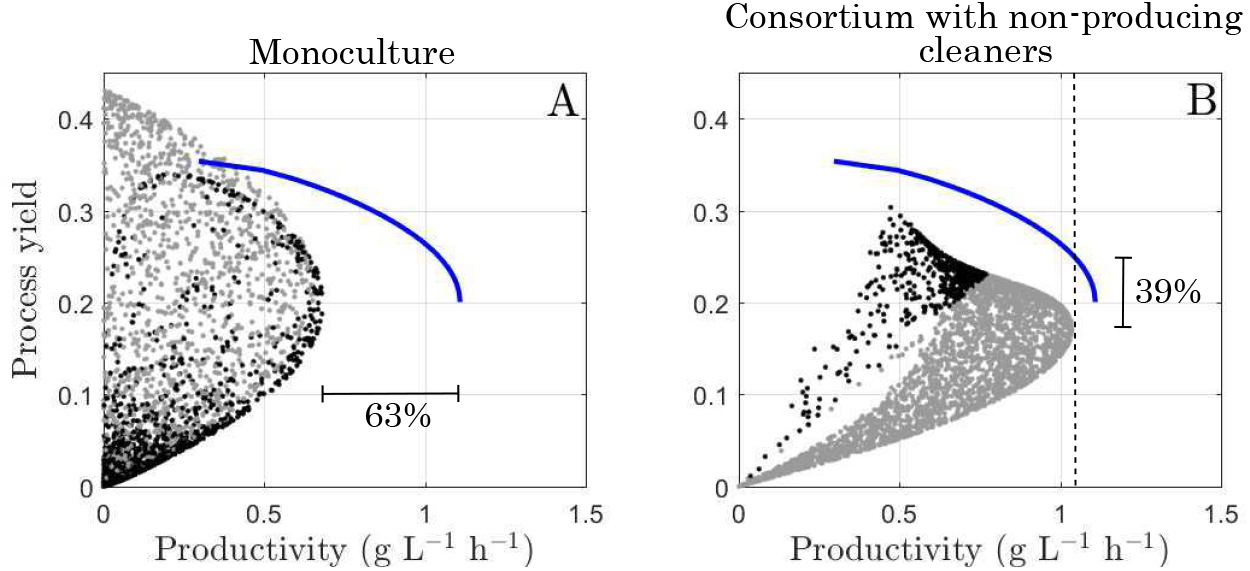
Scatter plots showing the performance of two alternative production methods, compared with Pareto-optimal performance of the consortium with producers and cleaners both synthezising the target protein. **(A)** Monoculture of producers: scatter plot of 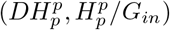, with 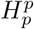 the protein concentration associated with the non-trivial equilibrium given by Proposition 3.1. Black points correspond to *D > D*^*a*^ and gray points correspond to *D* ≤ *D*^*a*^. The continuous line is the POF given by Figure 4. **(B)** Consortium of producers and non-producing cleaners: scatter plot of Φ with the protein yield *Y*_*h/c*_ fixed to 0. The scatter plot corresponds to Φ(*p*) with *v* ∈ Ω ∩ {*v*; *Y*_*h/c*_ = 0} and, as in panel A, the continuous line is the POF from Figure 4. Black points correspond to *D < D*^*c*^ and gray points correspond to *D* ≥ *D*^*c*^.

### 4.3 Cleaners must produce proteins for a better performance

In contrast with the work of Mauri et al. (2020), where cleaners cannot produce proteins, model (8)-(9) accounts for the production of recombinant proteins by cleaners. The question is whether there is a significant difference in terms of productivity when cleaners can produce proteins. To answer this question, we compare the POF obtained in Figure 4 with a scatter plot of the feasible objective region Φ(Ω) restricted to *Y*_*h/c*_ = 0 (see Figure 6B). In other words, the scatter plot shows the different process yields and productivities that can be reached when cleaners are only dedicated to remove acetate. The first observation is that allowing cleaners to produce proteins increases the process yield by 39% for a productivity of 1.04 g L^−1^ h^−1^, which is the maximal productivity for *Y*_*h/c*_ = 0. This increase is mainly explained by the fact that, when cleaners produce proteins, the problem of carbon diversion is addressed, that is, acetate secreted by producers is not wasted but transformed into proteins by cleaners.

## 5 Discussion and conclusions

### 5.1 Characterization of coexistence

We have established necessary and sufficient conditions for coexistence in a microbial consortium to synthesize recombinant protein. This consortium comprises two *E. coli* strains, the producer and the cleaner. Producers grow primarily on glucose while cleaners grow primarily on acetate that is secreted by producers as a fermentation by-product.

Conditions for coexistence are determined from analyzing the equilibria that each population can reach when growing separately. An important necessary condition for coexistence is that the producer secretes acetate at equilibrium, otherwise glucose is the only limiting resource and coexistence is impossible because of the competitive exclusion principle (Hardin, 1960). Assuming that this necessary condition holds, one simple description of our main result is obtained from the dynamical properties of the non-trivial equilibria on the boundary (*i*.*e*., equilibria in which only one strain is present). If each non-trivial equilibrium on the boundary repels solutions such that both strains grow, then coexistence is possible (see Corollary 3.8).

Another convenient description of our main result follows from an invasion analysis (Chesson, 2000). In our case, an invasion analysis amounts to choosing the *E. coli* strains one at time as the invader. The invader density is set to zero and the other strain, the so-called resident, is allowed to reach equilibrium. If the invader is then allowed to grown and attains a positive growth rate, we say that the invader has succeeded. If both strains can succeed as an invader, then they are said to coexist. The conditions for coexistence in our main result, Theorem 3.6, ensure the success of each strain as invaders. For example, in case (b) of the theorem, each strain reaches a non-trivial equilibrium when playing the role of resident, corresponding to the satisfaction of the two inequalities defining the necessary and sufficient conditions for this case.

The predominant inter-species interaction during coexistence can be competition, proto-cooperation, or syntrophy (Roell et al., 2019). From Theorem 3.6, we distinguish two cases in which coexistence is possible. In case (b), cleaners can survive growing separately, that is, producers are not necessary for the survival of cleaners. In this case, when both strains are grown together, producers may have either a positive or negative effect on cleaners. If producers secrete acetate at a low rate, then both populations strongly compete for glucose, which results in low growth of cleaners. Note that coexistence is possible because of acetate secretion, but the culture is dominated by competition. When the culture is operated at conditions close to washout of the monoculture of cleaner, then producers enhance cleaner growth by supplying acetate. This is known as protocooperation (Roell et al., 2019). In case (c) of Theorem 3.6, cleaners cannot survive growing individually. Coexistence is possible due to a syntrophic relationship in which the survival of cleaners critically depends on the secretion of acetate by producers.

The range of conditions allowing coexistence is affected by the metabolic burden of recombinant protein production (Kurland and Dong, 1996; Wu et al., 2016), which is in our case modulated by the values of the production yield parameters *Y*_*h/p*_ and *Y*_*h/c*_. A necessary condition for coexistence is that the monoculture of producers secretes acetate at equilibrium. Since the growth rate at equilibrium equals the dilution rate, and the secretion of acetate is directly related to the growth rate, this necessary condition can be put in terms of the dilution rate. Indeed, as shown in Proposition 3.1, the dilution rate must be higher than a threshold rate denoted by *D*^*a*^ and lower than a critical rate denoted *D*^*p*^. It can be proven from the definitions of *D*^*p*^ and *D*^*a*^ in Proposition 3.1 that

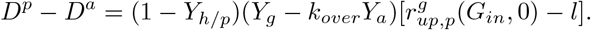

This shows how increasing *Y*_*h/p*_ decreases the range of dilution rates for which coexistence is possible.

### 5.2 Optimization: Coexistence, distribution of labor, and syntrophy

Using our results on the coexistence of the consortium, we numerically solved the multiobjective optimization problem (MOP) of maximizing both the process yield and productivity at the coexistence equilibrium. The performance of the consortium depends on different operational and strain design parameters. We chose as decision variables the dilution rate, a funtamental parameter of bioreactor operation, and product yields of the two strains, key parameters in the design and bioengineering of the synthetic community.

A first important question is whether the coexistence equilibrium is advantageous from a biotechnological point of view. As shown in Figure 6A, the *E. coli* consortium can reach a higher productivity than the monoculture of producers. However, the monoculture can reach a higher process yield. This is because of the inherent inefficiency associated with the loss of carbon through the intermediate resource (acetate). Indeed, the highest process yields for the monoculture are obtained in the absence of overflow metabolism (gray points in Figure 6A). One important observation is that all the elements of the Pareto optimal front are obtained when coexistence requires syntrophy, that is, when cleaners need the producers to survive. While this property holds in the case of the parameter values for the *E. coli* consortium (Mauri et al., 2020), we were not able to prove if it is a structural property of the system.

Division of labor refers to the execution of different tasks by different species in a consortium that are specialized for their respective tasks (Roell et al., 2019) and is often proposed as an effective design strategy in synthetic biology (Tsoi et al., 2018). Accordingly, Mauri et al. (2020) assumed that the production of proteins is the task of only one species, the producers. Another example is provided by Liu et al. (2018) who consider a syntrophic *E. coli* co-culture where only one strain synthesizes salidroside, the product of interest. Our results suggest that sharing the synthesis of the product of interest among the different species or strains enables higher yield and productivity. If only producers synthesize proteins, then the carbon lost through overflow metabolism is not eventually transformed into proteins. Alternatively, if only cleaners synthesize proteins, then carbon is inherently lost to producers through glucose competition. Distributing the task of producing proteins over the two strains reduces the loss of resources at the expense of increasing the metabolic burden of both populations. As a consequence, the trade-off between yield and productivity must be optimized simultaneously for both strains, leading to a global multi-level optimization problem. The example illustrates that, while division of labor may have advantages in terms of modularity and conceptual simplicity, it is not always optimal.

## Acknowledgements

This work was supported by the INRIA (IPL CoSy) and by the ANR projects Maximic (ANR-11-LABX-0028-01) and Ctrl-AB (ANR-20-CE45-0014). Additional support for Carlos Martínez was provided by the European Union within ESIF in the framework of the Operational Programme ”Research, Development and Education” (CZ.02.2.69/0.0/ 0.0/18 053/0016982).

## A Proofs

In this appendix we present the proofs of all the statements in Section 3 and we also present a series of technical results needed for their proofs. Throughout the appendix, we assume that (10), (11), and (2) hold.

*Proof*. (of Proposition 3.1) First, note that if (11) holds, then 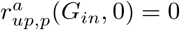 and 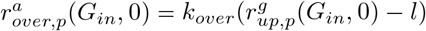. Consequently,

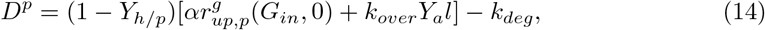

with *α* = *Y*_*g*_ − *k*_*over*_*Y*_*a*_ *>* 0 (see (10)). We also note that (see (11)):

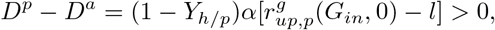

hence *D*^*p*^ *> D*^*a*^. Now, the steady states of (8) in absence of cleaners are given by the solution of the following system

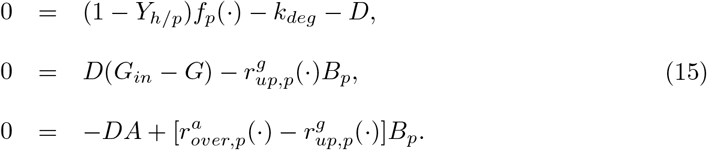

Following the same arguments of Proposition 1 by Martínez and Gouzé (2021), it is possible to show that

1. If *D* ≥ *D*^*a*^, then any solution of (15) satisfies 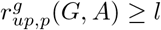.
2. If *D < D*^*a*^, then any solution of (15) satisfies *A* = 0 and 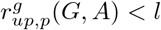.

If (1) holds, according to the property (7), we have that 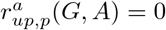. Consequently, (15) is equivalent to

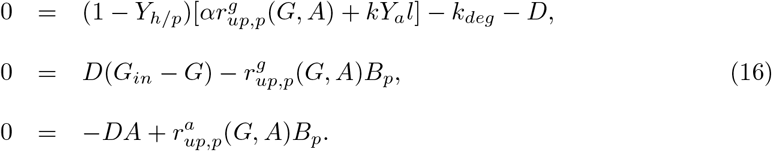

Again, following the proof of Proposition 1 by Martínez and Gouzé (2021), we have that

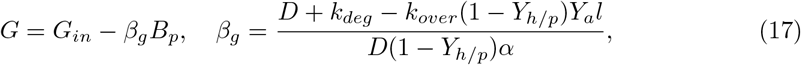

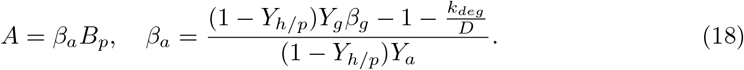

Replacing (17) and (18) in the first equation of (16), we obtain the following equation for *B*_*p*_:

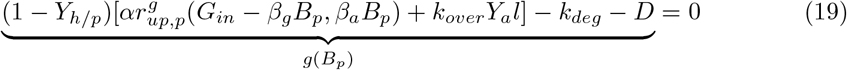

Since *g* is strictly decreasing, *g*(0) = *D*^*p*^ − *D*, and

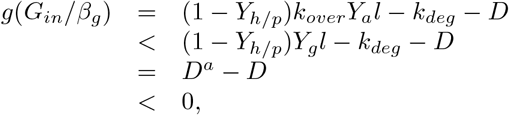

we conclude that (19) admits a unique solution 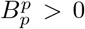 if *D < D*^*p*^, and has no positive solution if *D* ≥ *D*^*p*^. In particular, this proves (b).

If (2) holds, then *A* = 0 and 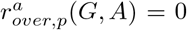. Replacing *A* and 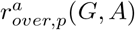 by 0 in (15), we obtain

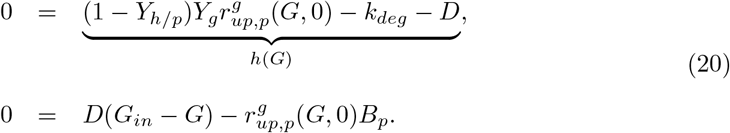

As in a classical chemostat model (note that *G* ↦ *h*(*G*) is strictly increasing), (20) admits a unique positive steady state if, and only if, *h*(*G*_*in*_) *>* 0. Since *D < D*_*a*_ and (11) holds, we have

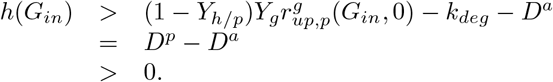

This completes the proof.

*Proof*. (of Proposition 3.4) Any equilibrium in absence of producers is a solution of

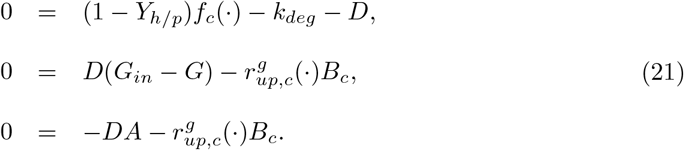

It is clear that *A* = 0. Consequently, we have

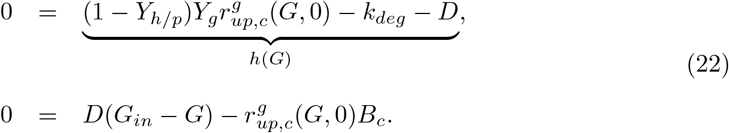

As in a classical chemostat model (Smith and Waltman, 1995), we have that (22) admits a solution, which is unique, if and only if *h*(*G*_*in*_) *>* 0. The rest of the proof follows from noting that *h*(*G*_*in*_) = *D*^*c*^ − *D*.

### Lemma A.1.

*Let D*^*a*^ *and D*^*p*^ *be defined by Proposition 3.1. If D* ≤ *D*^*a*^ *and* (1 − *Y*_*h/p*_) ≠ *β*(1 − *Y*_*h/c*_), *or D* ≥ *D*^*p*^, *then (8) has no coexistence equilibrium*.

*Proof*. If *D* ≤ *D*^*a*^, we claim that there is no coexistence equilibrium with acetate. Indeed, by contradiction, if 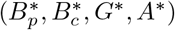 is a coexistence equilibrium of (8) with *A*\* *>* 0, from the fourth equation in (8) we obtain that

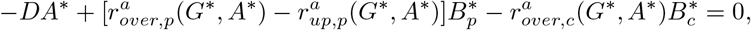

and therefore

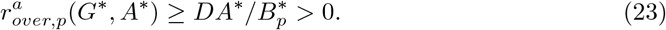

This implies that overflow metabolism occurs, hence, using (7), we have that there cannot be acetate uptake, 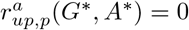. From the first equation in (8) and the definition of a coexistence equilibrium, we have that

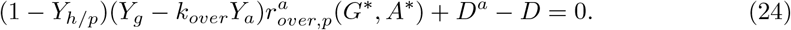

However, *D*^*a*^ ≥ *D* and 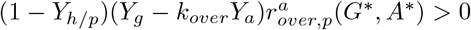, which contradicts (24). This proves that any coexistence equilibrium has no acetate. Therefore, any coexistence equilibrium is of the form 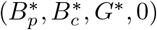 satisfying

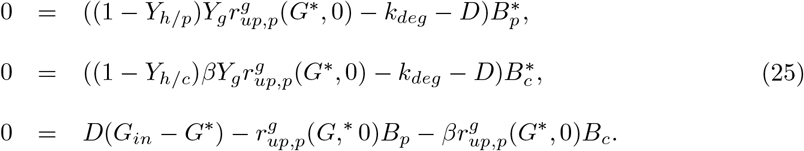

However, this implies that *B*_*p*_ = 0 or *B*_*c*_ = 0. Thus, there cannot be a coexistence equilbrium when *D* ≤ *D*^*a*^. This completes the proof.

### Lemma A.2.

*If (8) admits a coexistence equilibrium, say* 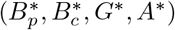, *then*

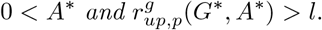

*Proof*. Let us assume that *D > D*^*a*^ and that (8) admits a coexistence equilibrium 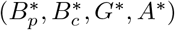. If *A*\* = 0, from the differential equation for *A* in (8), we obtain 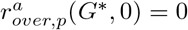 and consequently 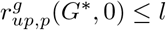. This implies that (see ODE for *B*_*p*_):

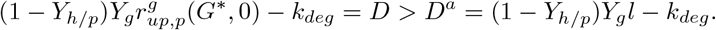

Hence, 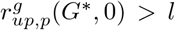, which is a contradiction. Therefore *A*\* must be positive, which implies that 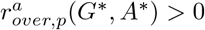 (same argument used to obtain (23)).

To continue studying the existence of a coexistence equilibrium, it is convenient to define two functions, *γ* and *ϕ*.

From Lemma A.1, coexistence is possible only when *D* ∈ (*D*^*a*^, *D*^*p*^), with *D*^*a*^ and *D*^*p*^ defined in 3.1. Under such conditions, overflow metabolism occurs at any equilibrium with producers (see Proposition 3.1 and Lemma A.2). Therefore, the glucose uptake rate 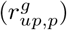 at any equilibrium with the presence of producers (*B*_*p*_ *>* 0) has the same value and is obtained from the following equation

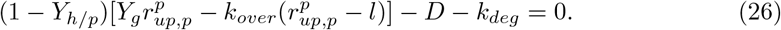

Let us define the function *γ* : (*D*^*a*^, *D*^*p*^) → ℝ_+_ by

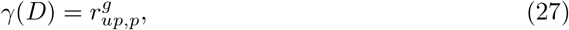

with 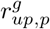 given by (27). It can be shown that

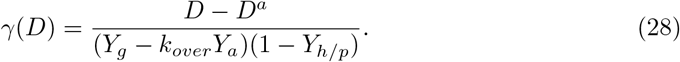

We also define the function *ϕ* : ℝ_+_ → ℝ by

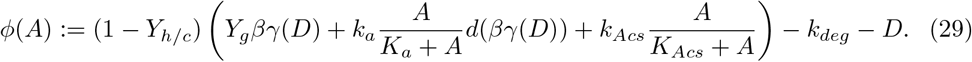

Note that if *A*\* is the acetate concentration at a coexistence equilibrium, then *ϕ* (*A*\*) = 0.

### Lemma A.3.

*Let D*^*a*^ *and D*^*p*^ *be defined by Proposition 3.1. If D* ∈ (*D*^*a*^, *D*^*p*^), *then (8) admits at most one coexistence equilibrium. Moreover, if a coexistence equilibrium exists, say* 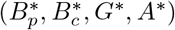, *then*

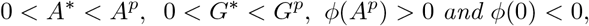

*where G*^*p*^ *and A*^*p*^ *are defined in Proposition 3.1*.

*Proof*. Let us assume that (8) admits a coexistence equilibrium 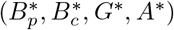. From the equation for *B*_*p*_ in (8), we have 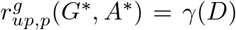, with *γ* defined by (27). From the definition of *G*^*p*^ and *A*^*p*^, we also have 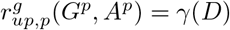. Similarly, we obtain that

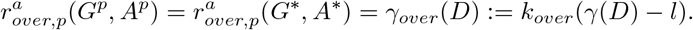

Now, we prove that *A*\* *< A*^*p*^. By contradiction, assume that *A*^*p*^ ≤ *A*\*. Using the monotonocity of *r*_*up,p*_ and the fact that 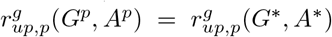, we obtain that *G*^*p*^ ≤ *G*\*. From the third equation in (8) we obtain that:

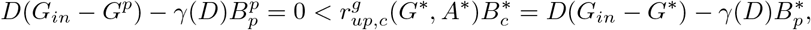

from where 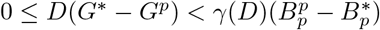 which implies

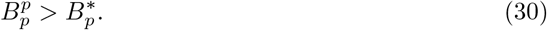

From the fourth equation in (8) we obtain that:

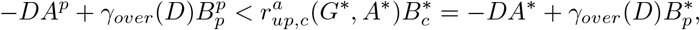

from where 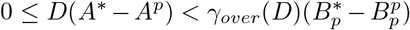 which implies 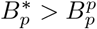. This contradicts (30). Then, *A*^*p*^ *> A*\* and consequently *G*^*p*^ *> G*\*. Now, from the equation for *B*_*c*_ in (8), we obtain *ϕ*(*A*\*) = 0, with *ϕ* defined by (29). Since *ϕ* is strictly increasing, we have the uniqueness of *A*^*c*^. The rest of the proof is straightforward.

### Lemma A.4.

*Let D*^*p*^ *and D*^*a*^ *be defined in Proposition 3.1 and assume that D*^*a*^ *< D < D*^*p*^. *Let D*^*c*^ *and G*^*c*^ *be defined by Proposition 3.1 and assume that D < D*^*c*^. *If* (1 − *Y*_*h/p*_)*f*_*p*_(*G*^*c*^, 0) − *k*_*deg*_ ≤ *D, then (8) has no coexistence equilibrium*.

*Proof*. Let us assume that (8) admits a coexistence equilibrium 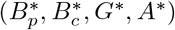. By definition we have

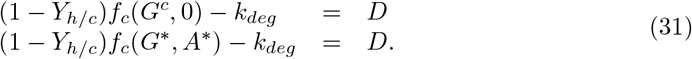

From Lemma A.3, we have that *A*\* *>* 0, hence 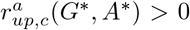. Recalling the definition of *f*_*c*_, from (31) we have that

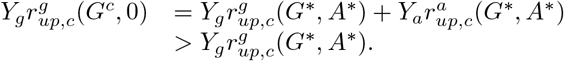

Consequently 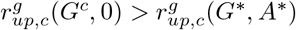. Since 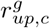 is proportional to 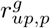, we have that 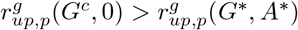. Now it is not difficult to show that

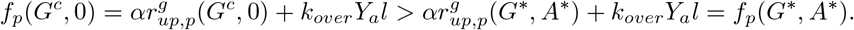

Now, if (1 − *Y*_*h/p*_)*f*_*p*_(*G*^*c*^, 0)− *k*_*deg*_ ≤ *D*, then (1 − *Y*_*h/p*_)*f*_*p*_(*G*\*, *A*\*)− *k*_*deg*_ *< D* which contradicts the definition of the coexistence equilibrium.

### Lemma A.5.

*Let D*^*p*^, *D*^*a*^, *and A*^*p*^ *be given by Proposition 3.1. If D*^*a*^ *< D < D*^*p*^, *then (8) admits a coexistence equilibrium if and only if*

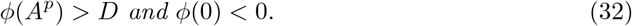

*Proof*. From Lemma A.3 we have that (32) is a necessary condition for the existence of a coexistence equilibrium. We prove here that it is also a sufficient condition. If (32) holds, since *ϕ* is strictly increasing, there exists a unique *A*\* ∈ (0, *A*^*p*^) such that *φ*(*A*\*) = 0. From the first equation in (8), we have *φ*(*G*) = 0 with *φ*(*G*) = (1 − *Y*_*h/p*_)*f*_*p*_(*G, A*\*) − *k*_*deg*_ − *D*. It is clear that *φ* is strictly increasing and that *φ*(0) = *−D−k*_*deg*_ *<* 0 and *φ*(*G*^*p*^) *>* (1 −*Y*_*h/p*_)*f*_*p*_(*G*^*p*^, *A*^*p*^) − *k*_*deg*_ −*D* = 0. Consequently, there is a unique *G*\* ∈ (0, *G*^*p*^) such that *φ*(*G*\*) = 0. It remains to prove the positiveness of the unique solution of the following linear system for (*B*_*p*_, *B*_*c*_):

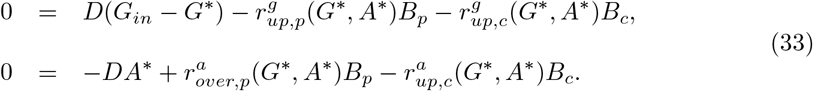

This system can be rewritten as

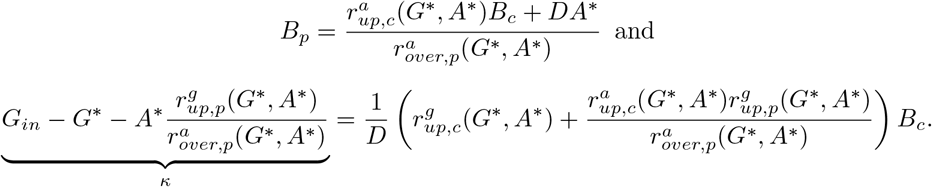

Since *A*^*p*^ *> A*\* and *G*^*p*^ *> G*\*, we have that

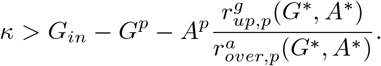

Finally, since 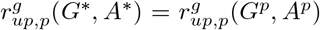, from (15) we conclude that *κ* = 0. Then (33) has a positive solution.

*Proof*. (of Theorem 3.6) Part (a) follows directly from Lemma A.2. Let us assume that *D*^*a*^ *< D < D*^*p*^. Using Lemma A.5, the proof of parts (b) and (c) follows from proving that

I. If *D < D*^*c*^ and (1 − *Y*_*h/p*_)*f*_*p*_(*G*^*c*^, 0) − *D* − *k*_*deg*_ *>* 0, then *ϕ*(0) *<* 0,
II. If *D* ≥ *D*^*c*^, then *ϕ*(0) *<* 0,
III. If (1 − *Y*_*h/c*_)*f*_*c*_(*G*^*p*^, *A*^*p*^) − *k*_*deg*_ − *D >* 0, then *ϕ*(*A*^*p*^) *>* 0.

For (I), from the hypotheses, we have that (1 − *Y*_*h/p*_)*f*_*p*_(*G*^*c*^, 0) − *k*_*deg*_ *> D*^*a*^. Using the definition of *D*^*a*^ we obtain *Y*_*g*_*l < f*_*p*_(*G*^*c*^, 0), and hence

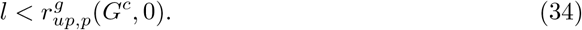

Now, from the definition of *G*^*p*^ and *A*^*p*^ (see ODE for *B*_*p*_), and from the hypotheses, we have that

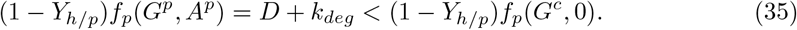

Since 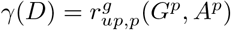 and 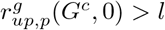, from (35) we conclude that

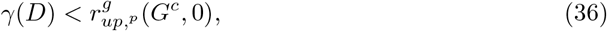

From the definition of *G*^*c*^ and (36) we obtain that

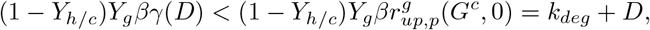

which implies *ϕ*(0) *<* 0.

For (II), we note that

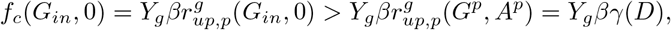

which implies

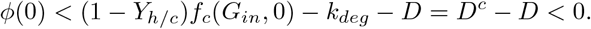

Finally, for (III), it is straightforward to verify that

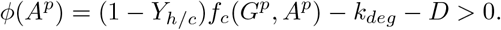

This completes the proof.

The following lemma, shows that the equilibrium with producers given by Proposition 3.1 is locally stable when seen as an equilibrium of (8) without cleaners.

### Lemma A.6.

*Assume that D > D*^*a*^. *Let* 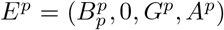 *be the equilibrium given by Proposition 3.1. Then* 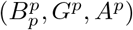 *is locally stable with respect to the following system:*

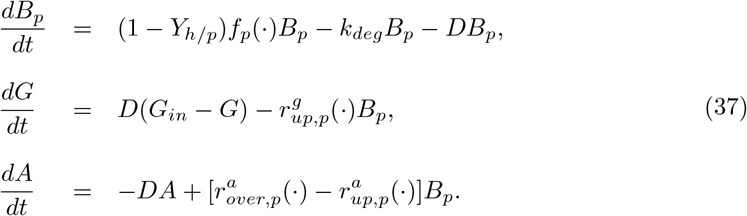

*Proof*. The proof follows the same idea of the proof of Proposition 2 by Martínez and Gouzé (2021). Along the proof, we will write *r* instead of 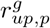. According to Proposition 3.1, 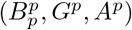 is in the region {(*B*_*p*_, *G, A*) ; *r*(*G, A*) ≥ *l*}. Therefore, using the definition of 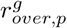 system and the property (7), we can study the local stability of 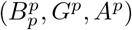 in the following system

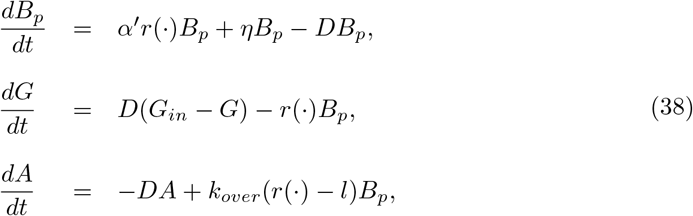

with *α*′ = (1−*Y*_*h/p*_)(*Y*_*g*_ −*k*_*over*_*Y*_*a*_) and *η* = (1−*Y*_*h/p*_)*k*_*over*_*Y*_*a*_*l* −*k*_*deg*_. The change of variables *U* = *B*_*p*_ + *α* ′*G* and *W* = *B*_*p*_ + (1 − *Y*_*h/p*_)*Y*_*g*_*G* + (1 − *Y*_*h/p*_)*Y*_*a*_*A* leads (38) to

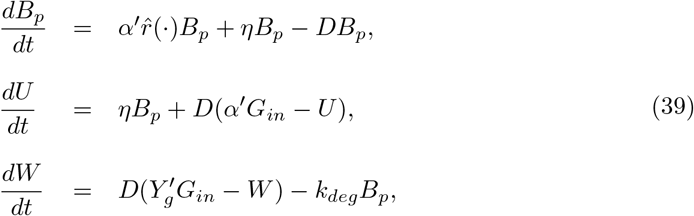

with

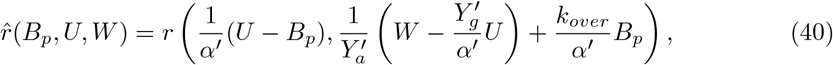

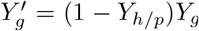, and 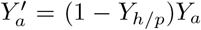. The characteristic polynomial associated with the Jacobian matrix of (39) is given by

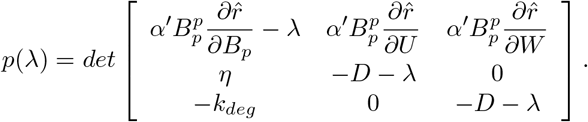

It can be shown that *p*(*λ*) can be written as

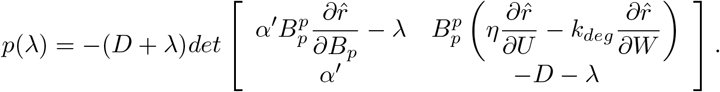

Thus, one root of *p* is −*D* and the other two roots are the eigenvalues of the matrix

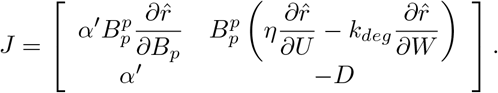

We note that:

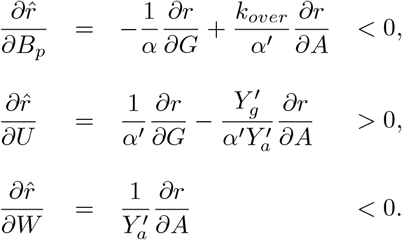

Using the fact that 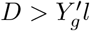, we have

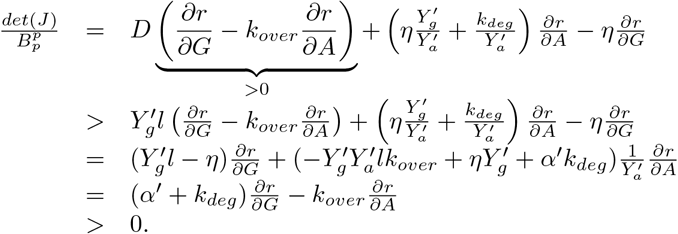

Therefore *det*(*J*) *>* 0. Now, it is clear that *Tr*(*J*) *<* 0, hence the eigenvalues of *J* have negative real part, and all the roots of *p* have negative real part. This completes the proof.

### Lemma A.7.

*Assume that D*^*a*^ *< D < D*^*p*^ *and let E*^*p*^ *be given by Proposition 3.1. Then the Jacobian matrix associated with (8) evaluated at E*^*p*^ *has three eigenvalues with negative real part, and one eigenvalue is given by:*

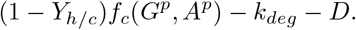

*Proof*. The evolution of *B*_*p*_, *B*_*c*_, *G*, and *A* is given by

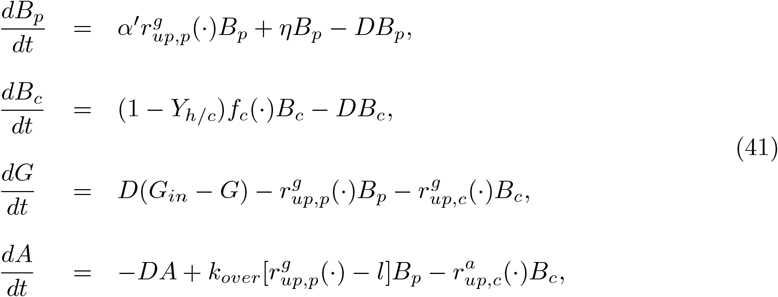

with *α*′ = (1 − *Y*_*h/p*_)(*Y*_*g*_ − *k*_*over*_*Y*_*a*_) and *η* = (1 − *Y*_*h/p*_)*k*_*over*_*Y*_*a*_*l* + *k*_*deg*_. The Jacobian matrix, *J*, associated with (41) evaluated at *E*^*p*^ takes the following form

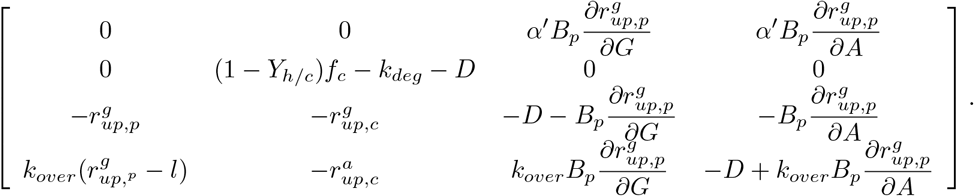

Using the properties of the determinant, we have that one eigenvalue is (1−*Y*_*h/c*_)*f*_*c*_(*G*^*p*^, *A*^*p*^)− *k*_*deg*_ − *D*, while the other three eigenvalues are exactly the same as those of the Jacobian matrix associated with (37) and evaluated at 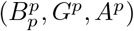. From Lemma A.6, we conclude that these three eigenvalues are negative. Thus, *E*^*p*^ is unstable and hyperbolic, if and only if, (1 − *Y*_*h/c*_)*f*_*c*_(*G*^*p*^, *A*^*p*^) − *k*_*deg*_ − *D <* 0.

### Lemma A.8.

*Assume that D < D*_*c*_ *and let E*^*c*^ *be the equilibrium given by Proposition 3.4. Then the Jacobian matrix associated with (8) evaluated at E*^*c*^ *has three eigenvalues with negative real part, and one eigenvalue given by*

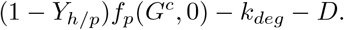

*Proof*. The Jacobian matrix of (8) evaluated at *E*^*c*^ is given by

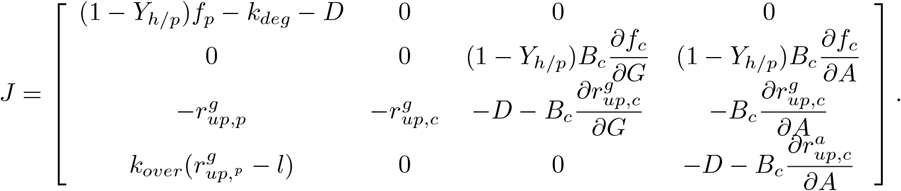

The characteristic polynomial associated with *J* takes the following form

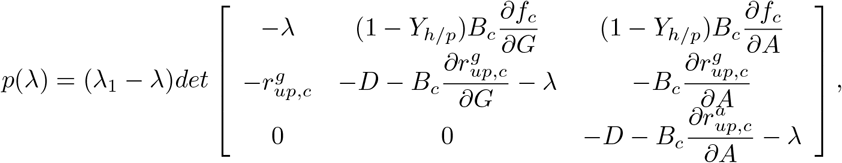

with *λ*_1_ = (1 − *Y*_*h/p*_)*f*_*p*_(*G*^*c*^, 0) − *k*_*deg*_ − *D*. Then we can write

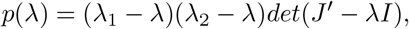

with 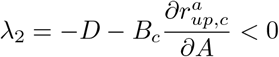 and

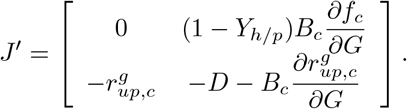

Since *Tr*(*J*′) *<* 0 and *det*(*J*′) *>* 0, *J*′ has two negative eigenvalues. Thus, *E*^*c*^ is hyperbolic and unstable if and only if *λ*_1_ *>* 0.

*Proof*. (of Lemma 3.10) Note that the variables 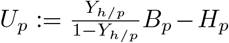 *and* 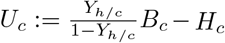 satisfy the following differential equations:

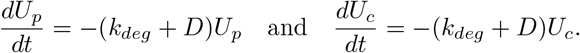

We conclude the proof noting that *U*_*p*_ and *U*_*c*_ asymptotically approach zero.

## B Algorithm to find the coexistence equilibrium

Let *D*^*a*^ and *D*^*p*^ be given by Proposition 3.1. From now on, we assume that *D* ∈ (*D*^*a*^, *D*^*p*^), otherwise there is no coexistence equilibrium (see Theorem 3.6). The first step is to determine the equilibrium 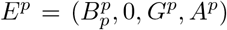 given by Proposition 3.1. The instructions on how to do so are dictated by the proof of Proposition 3.1. Indeed, 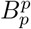 is obtained as the solution of

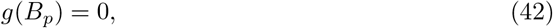

with *g* defined by (19). This equation has a unique solution on the interval [0, *G*_*in*_*/β*_*g*_], with *β*_*g*_ defined in (17). Moreover, *g*(0) *>* 0 and *g*(*G*_*in*_*/β*_*g*_) *<* 0, which provides an interval to look for the solution. Thus, equation (42) can be easily solved, for example, with the solver *fzero* in MATLAB. The values of *A*^*p*^ and *G*^*p*^ are obtained from

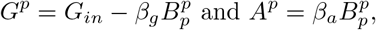

where *β*_*a*_ is defined in (18).

We also need the value of *G*^*c*^, the glucose concentration associated with the equilibrium *E*^*c*^ given by Proposition 3.4. Let *D*^*c*^ be given by Proposition 3.4. If *D < D*^*c*^, then *G*^*c*^ is the unique solution of (1 −*Y*_*h/c*_)*f*_*c*_(*G*, 0)− *k*_*deg*_ −*D* = 0. This equation is easily solved explicitly. If *D* ≥ *D*^*c*^, we will take *G*^*c*^ as *G*_*in*_. This is useful to distinguish the cases (b) and (c) in Theorem 3.6.

Now, to determine the coexistence equilibrium we use the following algorithm:

1. Determine *c*_1_ = (1 − *Y*_*h/p*_)*f*_*p*_(*G*^*p*^, *A*^*p*^) − *k*_*deg*_ − *D* and *c*_2_ = (1 − *Y*_*h/c*_)*f*_*c*_(*G*^*c*^, 0) − *k*_*deg*_ − *D*.
2. If *c*_1_ ≤ 0 or *c*_2_ ≤ 0, then there is no coexistence equilibrium. The algorithm ends. However, if *c*_1_ and *c*_2_ are positive, go to the next step.
3. Find *A*\* ∈ [0, *A*^*p*^] as the unique solution of *ϕ*(*A*) = 0, with *ϕ* defined by (29). Note that *ϕ*(0) *<* 0 and *ϕ*(*A*^*p*^) *>* 0.
4. Find *G*\* ∈ [0, *G*_*in*_] as the unique solution of *φ*(*G*) = 0 with *φ* (*G*) := (1−*Y*_*h/p*_)*f*_*p*_(*G, A*\*)− *k*_*deg*_ − *D*. Note that *φ* (0) *<* 0 and *φ* (*G*_*in*_) *>* 0.
5. Find 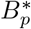 and 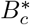 as the unique solution of the following linear system:

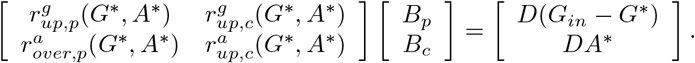

## C Algorithm to solve the MOP

Problem (13) is solved numerically with the interior point algorithm implemented in the toolbox *fmincon* of MATLAB (Byrd et al., 1999). To use *fmincon*, the objective function must be continuous on the feasible region which must be a closed set. In the following, we rewrite (13) in the standard form to use *fmincon*.

We recall the set Ω defined in Section 4. Consider the compact set 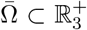 which can be described such that each element *p* = (*Y*_*h/p*_, *Y*_*h/c*_, *D*) on 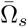 satisfies

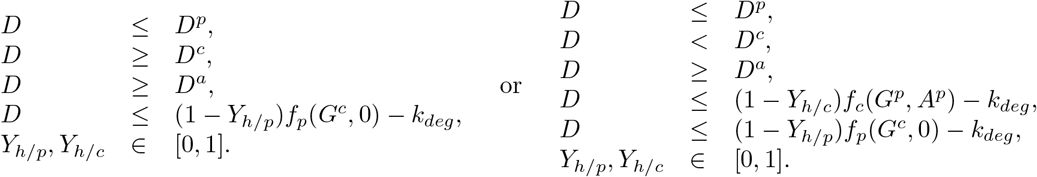

where *D*^*a*^, *D*^*p*^, *G*^*p*^, and *A*^*p*^ are given by Proposition 3.1, and *D*^*c*^ and *A*^*c*^ are given by Proposition 3.4. To extend the definition of Φ (defined in Section 4) on the boundary of 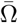 note that:

- If *D* ≥ *D*^*c*^, then cleaners can never survive (Proposition 3.4). Hence, we expect that solutions of (8) converges either to 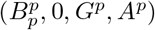 or to (0, 0, *G*_*in*_, 0).
- There is no element of the boundary satisfying *D* = *D*^*a*^. Indeed, as in the proof of Theorem 3.6, it can be proven that if *D < D*^*c*^, (34) holds. Now, in 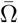, we have that *D* ≤ (1 − *Y*_*h/p*_)*f*_*p*_(*G*^*c*^, 0)− *k*_*deg*_. Combining this equation with (34), it is straightforward to show that *D > D*^*a*^.
- If *D < D*^*c*^ and *D* = *D*^*p*^, then there is no equilibrium with producers.
- If *D < D*^*c*^, *D < D*^*p*^ and *D* = (1 − *Y*_*h/p*_)*f*_*p*_(*G*^*c*^, 0) − *k*_*deg*_, then the equilibrium with producers is unstable.

Then we define

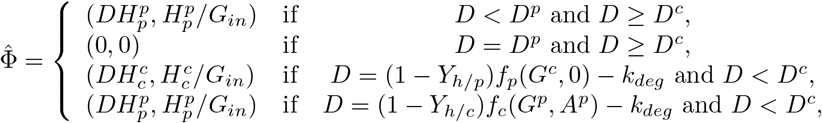

where 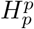 and 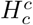 are the protein concentration associated with producers and cleaners, respectively. Then, we solve numerically the following problem instead of (13):

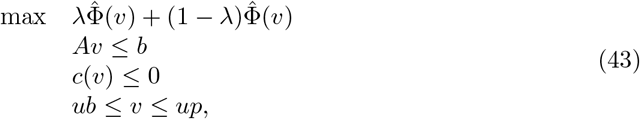

where

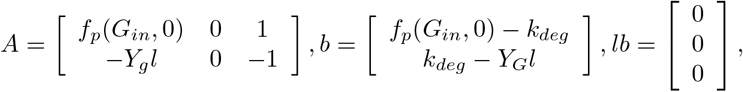

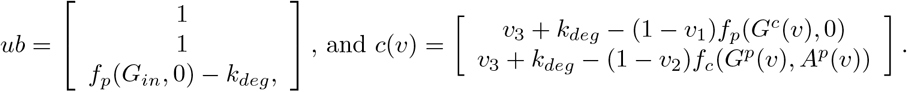

*G*^*c*^(*v*) is defined as *G*_*in*_ when *v*_3_ ≥ *D*^*c*^.

## D Robust choice of the down-regulation function

Let us assume that in (8) the down-regulation function *d* is replaced by 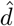, defined in (5). We then run long-term simulations of (8)-(9) until a steady state is detected (which is always the case).^2^ Then, we evaluate the the process yield and the productivity of the system at the time at which steady state has been reached. Figure 7 shows the result of this experiment for 1000 different values of (*Y*_*h/p*_, *Y*_*h/c*_, *D*). As we can see, the POF obtained when *d* is given by (6) represents a good approximation of the POF when *d* is given by (5). This shows that the choice of *d* in this paper is adequate to study the model proposed by Mauri et al. (2020).

**Figure 7:**
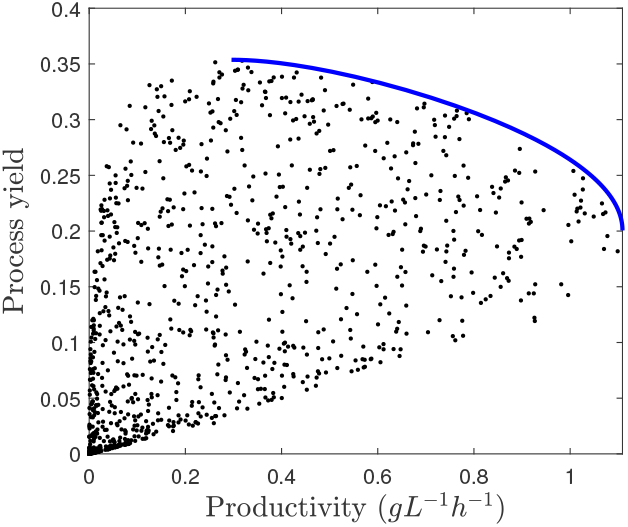
Scatter plot of Φ using the steady states of model (8)-(9) with *d* replaced by (5). The continuous line is the POF obtained in Figure 4

## Footnotes

1 This can be seen by noting that *d* is decreasing and that *d*(*l*) = 0.

2 Let *ξ* be a state variable of the model. A steady-state is detected when |*ξ*(*t* + Δ*t*) − *ξ*(*t*)| *< δ*. In our simulations we choose *δ* = 10^−6^ g L^−1^ and Δ*t* = 10 days.

